# High-Coverage Jomon Genomes Provide Insights into Population Structure and Genetic Traits of Ancient Japanese Hunter-Gatherers

**DOI:** 10.64898/2025.12.14.694187

**Authors:** Masahiko Kato, Fuzuki Mizuno, Yasuhiro Taniguchi, Osamu Kondo, Masami Matsushita, Takayuki Matsushita, Aiko Saso, Minoru Yoneda, Kunihiko Kurosaki, Li Wang, Koji Ishiya, Saki Aoto, Yusuke Watanabe, Mariko Isshiki, Izumi Naka, Jonghyun Kim, Shintaroh Ueda, Jun Ohashi

## Abstract

We analyzed eight high-coverage Jomon genomes spanning 9,000–2,800 years ago across the Japanese archipelago. Population genomic analyses revealed substantial genetic homogeneity across time and space reflecting post-glacial isolation, yet temporal stratification emerged: Initial Jomon (Iyai) individuals formed the earliest branch with weaker affinities to present-day Japanese. Using genomes from Todoroki and Chidorikubo sites, we estimated Jomon ancestry in mainland Japanese at ∼20%, substantially higher than previous estimates (∼13%). Polygenic score analyses, based on the direct analysis of Jomon genomes, provided the genetic evidence for their phenotypic predispositions, including elevated BMI and metabolic trait scores, reduced immune-related scores, and larger cardiac dimensions, patterns consistent with hunter-gatherer adaptations. These ancestral genetic influences, including elevated risks for obesity and dyslipidemia, persist in present-day Japanese. Our findings demonstrate that Jomon populations maintained genetic continuity despite ecological diversity, while their greater-than-expected contribution to present-day Japanese reshapes population history models and reveals how ancient adaptations continue influencing contemporary disease susceptibility.

## 1. INTRODUCTION

The dispersal of anatomically modern humans into East Asia began more than 40,000 years ago (Bae et al., 2017), but the demographic processes that shaped the genetic diversity of present-day populations remain incompletely understood. The Japanese archipelago, located at the eastern edge of the Eurasian continent, provides a unique setting for investigating human population history due to its geographic isolation and rich archaeological record. During the Last Glacial Maximum and subsequent deglaciation, rising sea levels between 20,000 and 15,000 years ago separated the Japanese landmass from the continent (Lambeck et al., 2014; Guedes et al., 2016). This isolation created an insular environment that profoundly influenced the genetic trajectory of its inhabitants, fostering the development of the Jomon culture—one of the world’s earliest pottery-using societies whose genetic components persist in present-day Japanese populations.

The Jomon period, spanning approximately 16,500 to 2,900 years before present(Habu, 2004), represents a striking example of long-term cultural and genetic continuity. Archaeological evidence indicates that Jomon people maintained a hunter-gatherer-fisher lifestyle across diverse ecological zones. They also developed pottery technology, sedentary settlements, and complex ritual practices (Habu, 2004; Craig et al., 2013; Kawashima, 2010). The prevailing “dual-structure model” of Japanese population history, originally proposed on the basis of cranial metric analyses, posits that present-day mainland Japanese emerged through admixture between indigenous Jomon hunter-gatherers and continental migrants who introduced rice agriculture during the Yayoi period (Hanihara, 1991; Mizoguchi, 2013). In this study, the term “mainland Japanese” specifically denotes Japanese populations other than the Ainu and Ryukyuan groups. This hypothesis was first supported by genetic studies of classical haploid markers, such as mitochondrial DNA and the Y chromosome (Tanaka et al., 2004; Hammer et al., 2006), which provided only limited resolution compared to genome-wide data. Mitochondrial analyses revealed high frequencies of haplogroups N9b and M7a among Jomon individuals, rare in continental populations (Adachi et al., 2011; Kanzawa-Kiriyama et al., 2013), …while Y-chromosome variation in populations closely related to the Jomon indicates a predominance of the Japan-specific lineage D-M55 (≈D1a2) (Tajima et al., 2004; Hammer et al., 2006; Nonaka et al., 2007; Sato et al., 2014).

Recent advances in ancient DNA sequencing have transformed our understanding of Jomon population history and their contribution to present-day Japanese ancestry. The first partial nuclear genome analysis of a 3,000-year-old Jomon individual from Fukushima demonstrated that the Jomon represented a basal East Asian lineage, having diverged before the subsequent diversification of East Asian populations (Kanzawa-Kiriyama et al., 2017). This finding was further substantiated by high-coverage whole-genome sequencing of Late Jomon individuals from the Funadomari site in Hokkaido. These genomes revealed that the Jomon lineage separated from continental populations approximately 20,000–15,000 years ago, coinciding with the geographic isolation of the Japanese archipelago (Kanzawa-Kiriyama et al., 2019). These studies established the Jomon as a genetically distinct population that experienced long-term isolation with a small effective population size of approximately 1,000 individuals. Comprehensive genomic analysis of a 2,500-year-old Jomon woman from the Ikawazu site provided evidence that the Jomon were descendants of an early coastal migration route from Southeast Asia to East Asia. The results also suggest that they were direct descendants of the Upper Paleolithic people who arrived in Japan around 38,000 years ago (Gakuhari et al., 2020). These ancient genome studies also revealed that admixture between Jomon and mainland Asian populations occurred at different times across regions—approximately 1,700–2,200 years ago in mainland Japanese and more recently (17–25 generations ago) in Hokkaido Ainu. These findings highlight the temporal and geographic heterogeneity of the admixture process (Gakuhari et al., 2020). In addition, a recent re-analysis of genome-wide data indicated that the Initial Jomon individual from Shikoku formed an outgroup to later Jomon populations, suggesting regional turnover within the archipelago (Jeong et al., 2023; Adachi et al., 2021). Large-scale analyses of present-day Japanese populations have revealed regional variation in the proportion of Jomon ancestry across the Japanese archipelago (Jinam et al., 2021; Watanabe and Ohashi, 2023).

While these findings have reshaped our understanding of the Jomon people as one of the ancestral source populations of the Japanese, fundamental questions remain unresolved. To date, only two Jomon individuals—from the Funadomari (Kanzawa-Kiriyama et al., 2019) and Iyai (Ishiya et al., 2024) sites—have been sequenced at high coverage, and conclusions based on other lower-coverage genomes require further validation. In particular, the Funadomari genome has frequently been used as a reference for the Jomon ancestry component, yet it represents an individual from Rebun Island at the northern edge of the Japanese archipelago. Given the likelihood of substantial regional diversity within the Jomon population, relying heavily on a single genome from such a peripheral locality raises questions about its representativeness when assessing the contribution of Jomon ancestry to present-day Japanese. Moreover, the genetic characteristics of the Jomon people, particularly those related to complex traits, have not yet been investigated with sufficient consideration of inter-individual variation (Martin et al., 2019). This gap underscores the importance of obtaining additional high-coverage genomes to more fully elucidate the genetic diversity of the Jomon people.

Here, we present whole-genome sequences from six newly sequenced Jomon individuals, analyzed together with data from two previously reported individuals, spanning the Initial to Late Jomon periods (approximately 9,000–2,800 years before present) from geographically diverse sites across the Japanese archipelago. By integrating high-coverage ancient genomes with comprehensive population genetic analyses, we aim to: (1) characterize the genetic structure and diversity of Jomon populations across time and space; (2) quantify differential genetic affinities between various Jomon groups and present-day Japanese populations; (3) refine models of Japanese population history as an admixture between Jomon and continental migrant groups; and (4) investigate the genetic basis of complex traits. Our expanded sampling reveals previously unrecognized temporal stratification within the Jomon gene pool and demonstrates how ancient hunter-gatherer adaptations continue to influence phenotypic variation in contemporary populations, providing comprehensive new insights into the interplay between demographic history and polygenic determinants of complex traits.

## 2. MATERIALS AND METHODS

### 2.1 Ancient sample information

We analyzed whole-genome sequence data from eight Jomon individuals spanning the Initial to Late Jomon periods. Two individuals had been reported previously: F23 from the Funadomari site (Late Jomon, Hokkaido) (Kanzawa-Kiriyama et al., 2019) and IY1 from the Iyai site (Initial Jomon, Gunma) (Ishiya et al., 2024). We additionally sequenced six individuals: TO5 from the Todoroki site (Early Jomon, Kumamoto); IY4, IY15, and IY18 from Iyai (Initial Jomon, Gunma); MI3 from Mitsusawa (Middle Jomon, Kanagawa); and C3 from Chidorikubo (Middle Jomon, Tokyo). Radiocarbon dates for TO5, IY4, IY15, IY18, MI3, and C3 had been obtained previously; in this study, we use only the 14C ages, primarily for assigning archaeological periods, and the full set of radiocarbon dating results will be reported in a separate publication. This study was approved by the Ethics Committee of Toho University School of Medicine (A24062_A23103_A20110_A18099_A18056).

### 2.2 DNA extraction, library preparation, and sequencing

Sample preparation followed a previous study (Mizuno et al., 2021). Briefly, we used six Jomon individuals (IY4, IY15, IY18, TO5, C3, and MI3) for whole-genome analysis. DNA was extracted from the right petrous bone of Individuals IY4, T05, and M3, and from the left petrous bone of Individuals IY15, IY18, and C3. Next-generation sequencing (NGS) libraries were constructed following protocols described in our previous studies. Sampling targeted the dense region of the petrous bone surrounding the cochlea. The outer surface of each bone was removed with a sanding tool (Dremel, USA), and the cleaned inner portion was pulverized into fine powder using a bead mill (Multi-Beads Shocker MB601U; Yasui Kikai, Japan). The powder was decalcified in 0.5 M EDTA (pH 8.0) at 56 °C for 2 hours in a rotating oven; after centrifugation, the supernatant was recovered, and this decalcification step was repeated three times. DNA was then extracted from the EDTA supernatant using phenol:chloroform:isoamyl alcohol (25:24:1), followed by an additional chloroform wash. The aqueous phase was collected, concentrated to 200 μL with Amicon Ultra-15 centrifugal filters (Merck Millipore, Germany), and further purified using a MiniElute silica spin column (QIAGEN, Germany). From the extracted DNA, both double-stranded (DS) and single-stranded (SS) NGS libraries were prepared for Individuals IY4, IY15, IY18, T05, C3, and M3, followed by shotgun sequencing. All libraries were amplified and initially sequenced on the Illumina MiSeq platform using the 150-cycle v3 kit to evaluate library complexity, DNA preservation, and authenticity. Based on these assessments, high-coverage sequencing was subsequently performed on the Illumina NovaSeq 6000 platform (150 bp paired-end reads) at SRL Inc., Macrogen, or Genewiz.

### 2.3 Sequencing data processing

IY15 was sequenced using paired-end sequencing across five separate runs. Adapter trimming was performed using fastp version 0.23.4 with the -c and -3 options. Following quality control, sequence files derived from the same sample were concatenated into a single file for downstream analysis. The sequencing data for IY15 were processed using the Illumina DRAGEN Bio-IT Platform (Host Software Version 07.021.645.4.0.3, Bio-IT Processor Version 0×18101306). This platform employs a hardware-accelerated pipeline that integrates read mapping, local alignment, duplicate marking, and haplotype variant calling in accordance with GATK Best Practices recommendations.

We processed the raw sequencing reads from each library using fastp (ver. 0.20.0) (Chen et al., 2018), trimming adapter sequences, filtering low-quality bases, and discarding short reads (< 35 bp). After filtering, we subsequently mapped the retained reads to the human reference genome sequence (hg19) with bwa aln (ver. 1.9) (Li et al., 2009) using parameters optimized for ancient DNA (-l 1024 -n 0.01 -o 2). We then applied the Picard Toolkit (ver. 2.21.4) (Broad Institute, 2019) MarkDuplicates command to remove PCR and optical duplicates from each library. Next, we merged the duplicate-removed BAM files across libraries and filtered out poorly aligned reads (MAPQ < 30) using SAMtools (ver. 1.9) (Li et al., 2009).

To confirm the authenticity of ancient DNA, we assessed nucleotide misincorporation patterns and DNA fragmentation using MapDamage (ver. 2.0.8) (Jónsson et al., 2013) with aligned reads from non-repaired NGS libraries. In addition, we calculated DNA contamination rates based on definitive haplogroup sites in the mitochondrial genome using MitoSuite (ver. 1.0.9) (Ishiya et al., 2019).

Before variant calling, we applied BamUtil (ver. 1.0.15) (McKennna et al., 2010) to trim 2 bp from both ends of each read in the merged BAM file, thereby reducing spurious mismatches caused by ancient DNA damage. We then performed a single-nucleotide variant (SNV) calling on the trimmed and high-quality alignments using GATK HaplotypeCaller (ver. 3.8.1) (DePrist et al., 2011;Jun et al., 2015). To minimize false-positive calls, we excluded multiallelic sites with ≥ 3 alleles and removed heterozygous sites with an alternate allele frequency below 0.2. Finally, we retained the high-confidence variant calls in VCF format and used them for downstream population genomic analyses.

### 2.4 Sex identification

Sex of each individual was estimated from the distribution of sequencing reads mapping to the sex chromosomes. For each library, we calculated the number of reads aligned to chromosomes X and Y using the idxstats function in SAMtools v1.19 (Li et al., 2009). We then computed the RY index, defined as Y/(X+Y), and assigned genetic sex according to previously established thresholds (Skoglund et al., 2013). These results are summarized in Table S1.

For two individuals (MI3 and C3), the initial RY values based on the primary sequencing data fell in an intermediate range and did not allow an unambiguous sex assignment. To refine these cases, we generated additional double-stranded (DS) libraries from the same skeletal material, performed whole-genome in-solution enrichment, and sequenced the enriched libraries on an Illumina MiSeq instrument. We again quantified the numbers of reads mapping to the X and Y chromosomes from these enriched datasets. Both MI3 and C3 exhibited substantial X-chromosomal coverage but only trace levels of Y-chromosomal reads, a pattern consistent with a female karyotype. These supporting data are provided in Table S2 and were used to classify MI3 and C3 as genetically female in the Results.

### 2.5 Reference datasets and data processing

Reference populations were assembled from two primary sources. First, ancient genome data were obtained from the Allen Ancient DNA Resource (AADR) v54.1 (Mallick et al., 2024), extracting 330 East Asian individuals and 18 previously reported Jomon genomes genotyped at 973,002 SNPs on the 1240K panel (this dataset includes published ancient genomes from the Japanese archipelago and Korean peninsula). Second, present-day Japanese data were derived from a previously published dataset of 11,069 individuals sampled across all 47 prefectures, using existing allele frequency data from 50 randomly selected individuals per prefecture (Watanabe et al., 2021).

Data were merged using mergeit (EIGENSOFT v7.2.1) (Patterson et al., 2006). AADR samples were incorporated without additional filtering, as they had undergone standardized quality control. The merged dataset captured East Asian genetic diversity across temporal and geographic dimensions, enabling robust population genetic analyses.

### 2.6 Population genetic analyses

#### 2.6.1 Principal component analysis

A principal component analysis (PCA) was performed using smartpca v16000 (EIGENSOFT v7.2.1) (Patterson et al., 2006) on 973,002 SNPs from the 1240K panel. The analysis included present-day East Asian populations and ancient individuals from Japan and Korea. Ancient individuals were projected onto PC space calculated from present-day populations using lsqproject: YES.

#### 2.6.2 ADMIXTURE analysis

Population structure was assessed using ADMIXTURE v1.3.0 (Alexander et al., 2009). Prior to analysis, we filtered SNPs for minor allele frequency (--maf 0.01) and pruned for linkage disequilibrium (--indep-pairwise 200 25 0.2), retaining 122,381 SNPs. K values from 2 to 10 were tested in 10 independent runs each, with cross-validation (--cv=10) used to identify the optimal K. The lowest CV error (0.47379) was observed at K=4.

#### 2.6.3 *f*-statistics analysis

To quantify genetic relationships, *f*-statistics were calculated using ADMIXTOOLS2 (Patterson et al., 2012; Maier et al., 2023). Outgroup *f*_3_-statistics, *f*_3_(Mbuti; X, Y), were used to assess genetic similarity among Jomon individuals, Yayoi- and Kofun-period individuals, and present-day Japanese and Koreans. To test for differential Jomon affinity, we calculated *f*_4_(Mbuti, Test; Jomon1, Jomon2), where Test was defined as either present-day Japanese or present-day Koreans. Standard errors were estimated through block jackknife resampling (714 blocks for qpf3), implemented in each software package. Statistical significance for *f*-statistics was assessed using |Z| > 3, corresponding to *P*-value < 0.003. This conservative threshold controls for multiple testing without formal correction. Outgroup *f*_3_-statistics, *f*_3_(Mbuti; prefecture, Jomon), were calculated with our custom implementations to assess genetic affinities between Jomon individuals and the 47 prefectural populations of Japan.

Data processing used Python 3.10 with NumPy and pandas. Visualizations were created using matplotlib and seaborn, except where software provided integrated plotting. Analysis scripts and data are available upon request from the authors.

#### 2.6.4 Admixture modeling

Jomon ancestry proportions were estimated using *f*_4_-ratio tests in qpF4ratio v400 (Patterson et al., 2019). The model assumed present-day Japanese descended from admixture between Jomon and continental East Asian populations. Following previous studies (Adachi et al. 2021; Kanzawa-Kiriyama et al., 2019), Dai served as the outgroup population, Koreans as continental source, and various Jomon populations as indigenous sources. SNP counts varied between 418,449 and 601,334 across sample combinations.

#### 2.6.5 Phylogenetic analysis

Phylogenetic relationships were inferred using TreeMix v1.13 (Pickrell et al., 2012). The analysis included eight Jomon individuals, present-day East Asian populations, and ancient samples from Japan and Korea. Trees were rooted with Mbuti as the outgroup population and evaluated with 500 bootstrap replicates. We ran TreeMix without migration edges (-noss option) to establish the tree topology based on population splits. Drift parameters were estimated for each branch to quantify genetic divergence since population separation.

#### 2.6.6 Polygenic score analysis

Polygenic scores (PSs) were calculated using summary statistics from genome-wide association studies (GWAS) conducted in BioBank Japan (http://jenger.riken.jp/result). The set of traits analyzed encompassed anthropometric measures, metabolic indicators, cardiovascular health, kidney function, and immune-related traits (Akiyama et al., 2017; Kanai et al., 2018; Akiyama et al., 2019). Effect sizes were extracted for SNPs with *P* < 0.01. To ensure reliable diploid genotype calls across SNP sites and maximize the number of SNPs available for analysis, PS calculations were restricted to six high-coverage Jomon individuals: IY1, IY4, TO5, MI3, C3, and F23. For comparison, mean PS values of present-day Japanese were calculated on the basis of allele frequencies from the Japanese population in the 1000 Genomes Project (1000 Genomes Project Consortium, 2012). Complete trait lists, along with the corresponding GWAS references, are provided in Table S6.

## 3. RESULTS

### 3.1 Inferred genetic sex of Jomon individuals

We first inferred the genetic sex of the eight high-coverage Jomon individuals based on the ratio of Y chromosome reads to total sex chromosome reads (RY index), following the criteria of Skoglund et al. (2013) (Table S1). Three individuals (IY4, IY15, and IY18) were classified as male and three (IY1, TO5, and F23) as female. Two individuals (MI3 and C3) initially showed intermediate RY values and were provisionally designated as undetermined. To further evaluate MI3 and C3, we generated double-stranded (DS) libraries and performed whole-genome in-solution enrichment followed by MiSeq sequencing. Both individuals showed substantial X-chromosomal coverage but only trace levels of Y-chromosomal reads (Table S2), consistent with a female karyotype. We therefore classified both MI3 and C3 as genetically female.

Individual IY1 was identified as a 20–40-year-old female based on skeletal morphology (Kondo et al., 2018), and genomic sex determination likewise confirmed this assignment (Ishiya, under review). Individual IY18 was assessed morphologically as an older adolescent to young adult male, which was consistent with the genomic results. In contrast, individuals IY4 (12–18 years old) and IY15 (12–15 years old) were too young for reliable morphological sex estimation; however, genomic analysis indicated that both were male. Individual TO5 was identified as a mature adult female (40–50 years old) based on morphological analysis, and genomic sex determination again confirmed this assessment. For MI3, only the cranial vault and mandible were preserved. Although the individual was previously considered male based on mandibular morphology, our genomic analysis, including the targeted whole-genome enrichment data described above, identified the individual as female. Individual C3 was estimated to be approximately four years old based on dental development, but morphological sex estimation was not possible. Genomic analysis, supported by the enrichment data, revealed that the individual was female. For IY1 and F23, our genomic sex determinations were also concordant with the findings of previous genomic studies (Kanzawa-Kiriyama et al., 2019; Ishiya et al., 2024).

### 3.2 Genetic structure of Jomon individuals and their position among East Asian populations

To characterize the genetic structure of Jomon individuals in the context of East Asia, we analyzed principal component analysis (PCA) and ADMIXTURE analysis using genome-wide SNP data from eight high-coverage Jomon genomes and a set of present-day East Asian reference populations.

PCA revealed a distinct population structure, with all Jomon individuals forming a tight cluster that was clearly separated from present-day East Asian populations along PC1 (Figure 2). Despite representing different temporal phases—Initial (IY1, IY4, IY15, and IY18), Early (TO5), Middle (MI3 and C3), and Late Jomon (F23)—and covering a wide geographic range from Hokkaido to Kumamoto, all eight Jomon individuals with high-coverage genomes clustered closely together. This compact clustering contrasts with the broader dispersion observed among previously available low-coverage Jomon genomes, underscoring the importance of sequencing depth in resolving fine-scale population structure.

**Figure 1.**
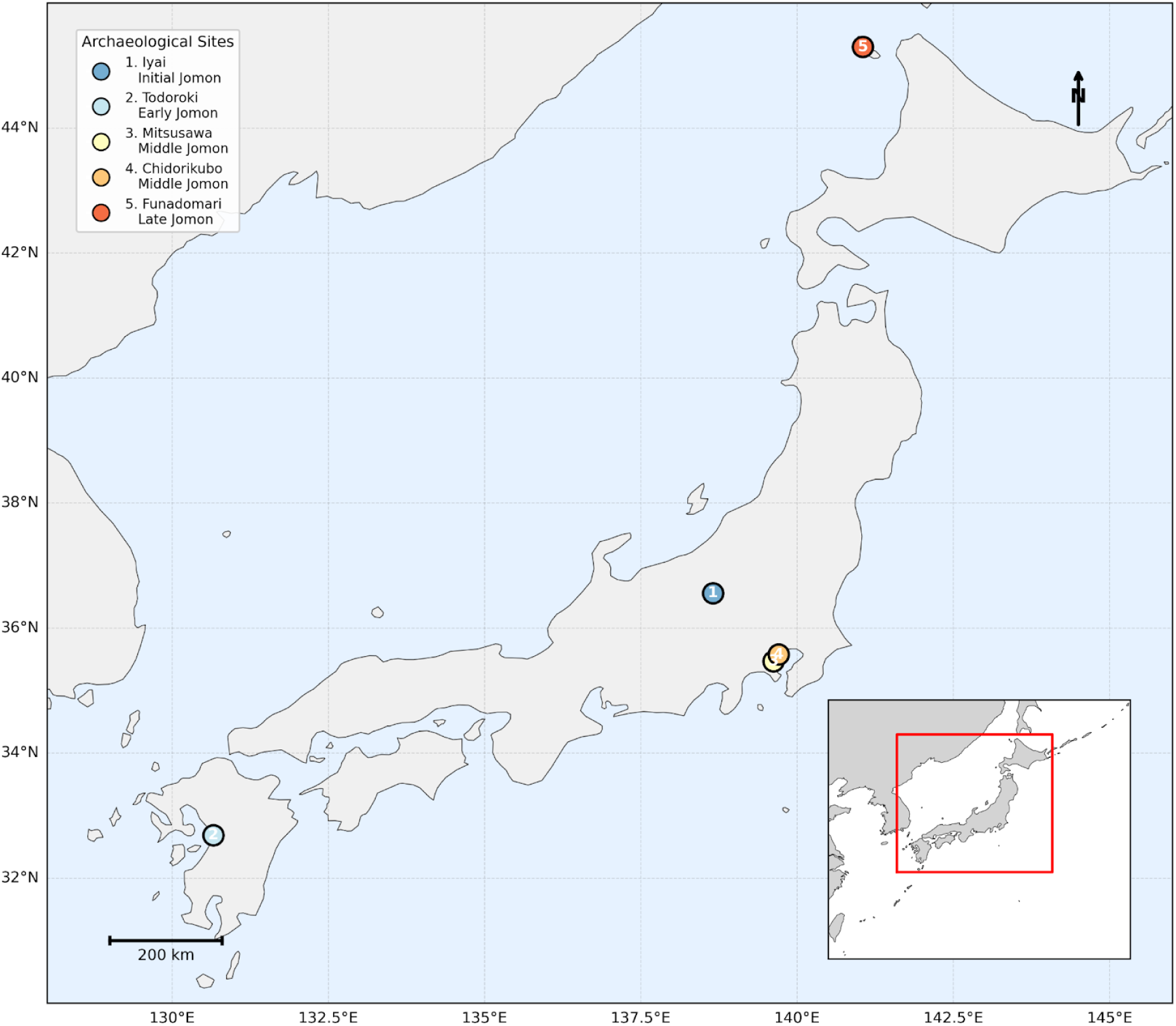
Geographic locations of the five Jomon archaeological sites analyzed in this study. Sites are numbered chronologically from oldest to youngest: (1) Iyai, Gunma Prefecture (Initial Jomon); (2) Todoroki, Kumamoto Prefecture (Early Jomon); (3) Mitsusawa, Kanagawa Prefecture (Middle Jomon); (4) Chidorikubo, Tokyo (Middle Jomon); and (5) Funadomari, Rebun Island, Hokkaido (Late Jomon).

**Figure 2.**
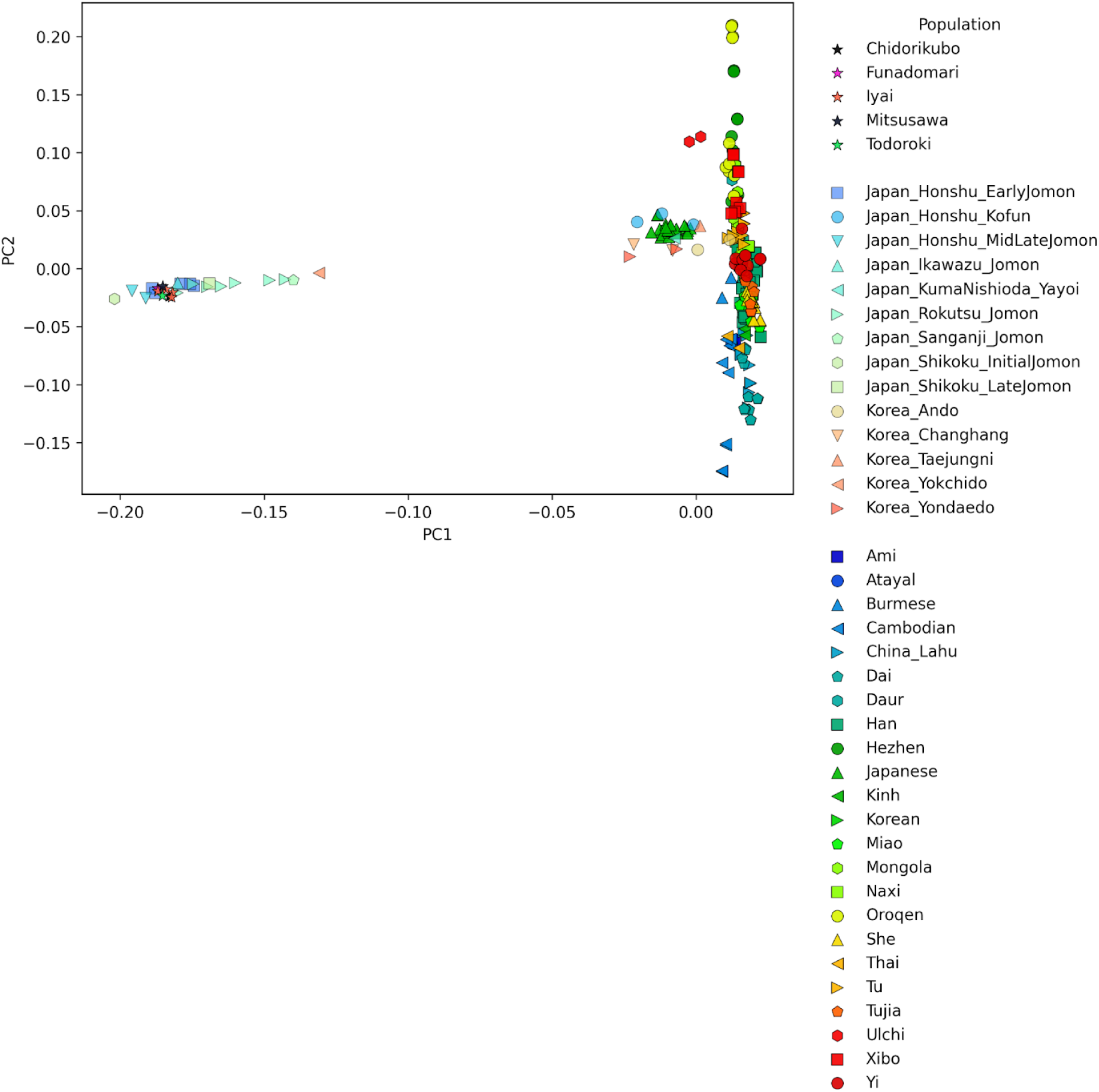
Principal component analysis of ancient and present-day East Asian populations. Principal component analysis was performed using 973,002 SNPs from the 1240K panel. Ancient individuals (including low-coverage Jomon samples from the AADR) are shown in light colors, whereas present-day populations are depicted in dark colors. PC1 and PC2 explain 6.865% and 3.342% of the total variance, respectively.

ADMIXTURE analysis (K = 4) revealed that all high-coverage Jomon individuals derived nearly all of their ancestry from a single genetic component (shown in blue), with only minimal contributions from continental East Asian sources (Figure 3). These findings indicate substantial genetic homogeneity across both temporal and geographic scales. Notably, although the Jomon period spanned more than 16,000 years, the genetic evidence strongly suggests that the Jomon population remained largely isolated from continental Eurasian groups throughout this time, with little to no gene flow into the population. Moreover, the genetic contribution of the Jomon people to present-day populations appears to have been largely confined to the Japanese, as the Jomon-derived component (shown in blue) is detected predominantly in present-day Japanese and is largely absent from other East Asian groups. Although a Jomon-derived component is detectable in populations of the Japanese archipelago after the Jomon period, it persists only at low levels. Most of the remaining ancestry reflects genetic contributions from continental migrants. Accordingly, continental migrants contributed the predominant share of ancestry to populations in the Japanese archipelago after the Jomon period.

**Figure 3.**
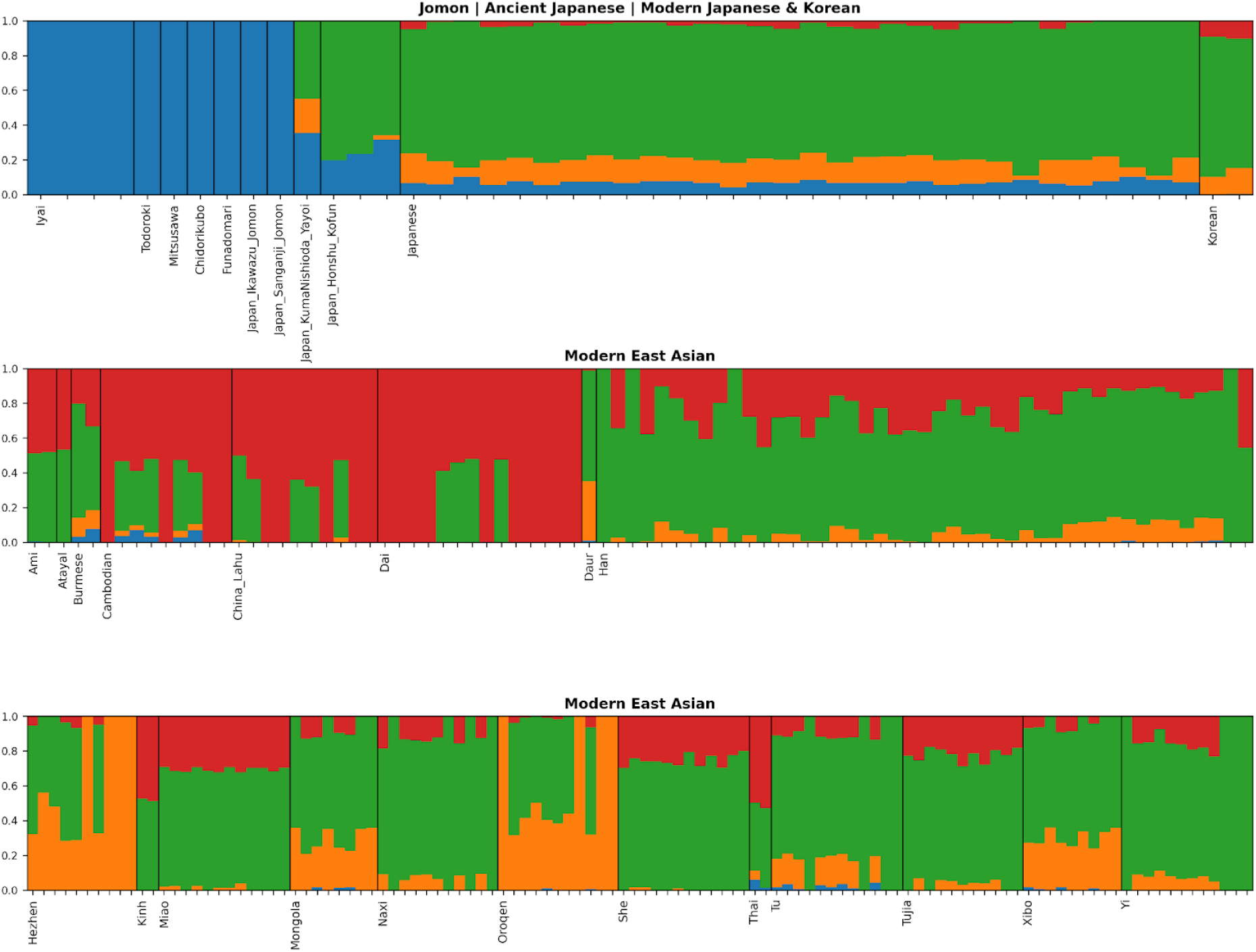
ADMIXTURE analysis supports genetic isolation of Jomon individuals. Unsupervised ADMIXTURE analysis at K = 4 (optimal based on the cross-validation error = 0.47379) is shown for eight high-coverage Jomon individuals; ancient and present-day Japanese and Korean individuals (top panel); and present-day East Asian populations (middle and bottom panels). Each vertical bar represents an individual, with colors denoting the corresponding ancestry components.

### 3.3 Genetic affinities within the Jomon and their divergence from later populations

To assess genetic similarity among Jomon individuals, Yayoi- and Kofun-period individuals, and present-day Japanese and Koreans, we calculated outgroup *f*_3_-statistics using Mbuti as the outgroup (Figure 4). We first evaluated pairwise genetic affinities among Jomon individuals from different time periods and geographic locations. The clear differences observed between the eight Jomon individuals highlighted in this study (IY1, IY4, IY15, IY18, TO5, C3, MI3, and F23) and the other seven Jomon individuals do not reflect population structure within the Jomon. Instead, they are attributable to differences in data quality, with the former represented by high-coverage genome data and the latter by low-coverage data. As with the PCA analyses, the use of low-coverage data in *f*-statistic calculations can introduce biases, which may obscure true genetic relationships and artificially inflate apparent differences among individuals.

**Figure 4.**
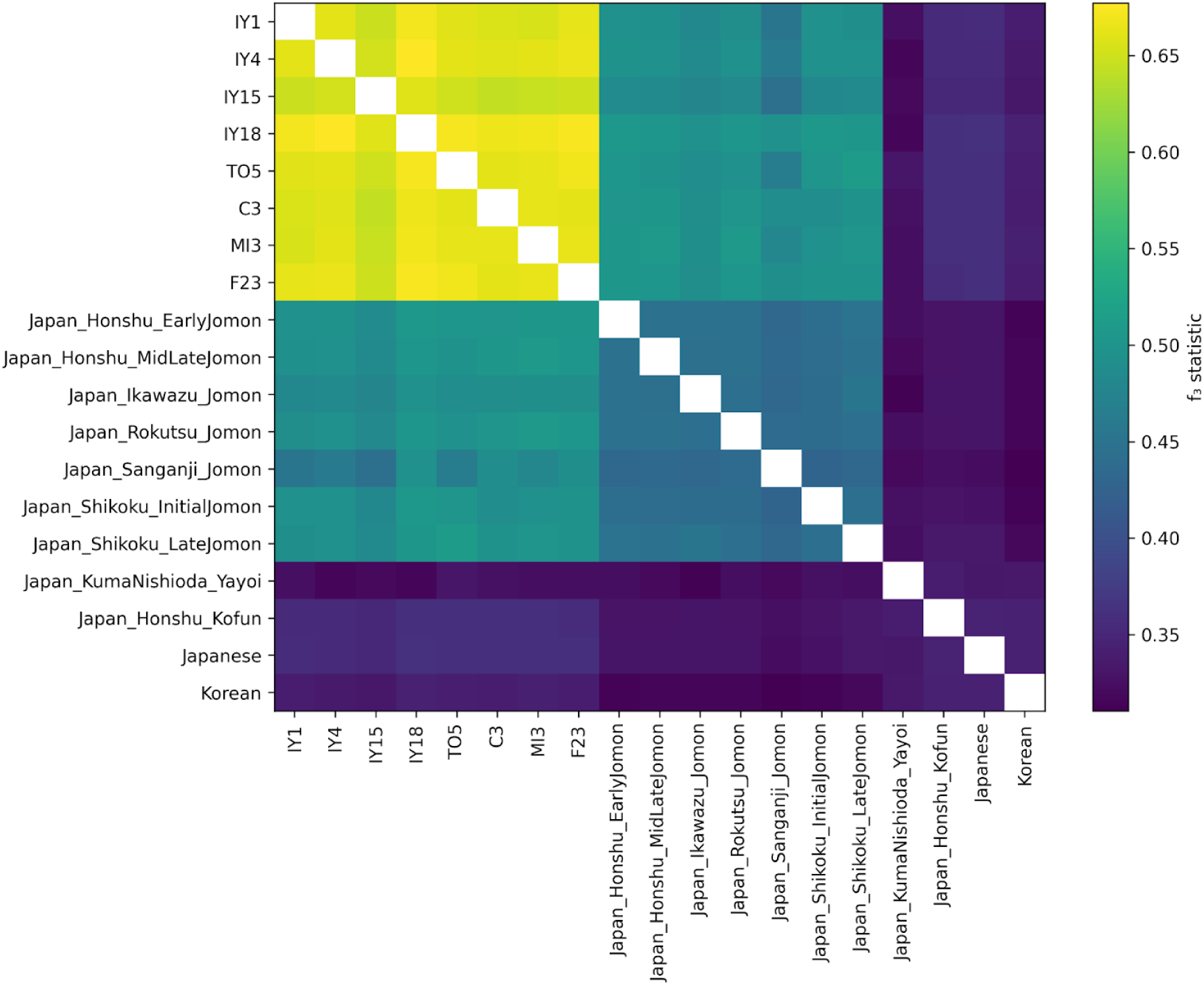
Pairwise genetic affinities among Jomon individuals based on outgroup *f*_3_ statistics. The heatmap shows *f*_3_ values in the form *f*_3_(Mbuti; X, Y), illustrating all pairwise comparisons among Jomon individuals, Yayoi- and Kofun-period individuals, and present-day Japanese and Koreans (multiple individuals each). Higher values (yellow) indicate greater shared genetic drift. All *f*_3_ values presented in this figure were significantly greater than zero (Z > 3).

The *f*_3_ values between Jomon and post-Jomon populations in the Japanese archipelago were substantially lower (<0.4), not only with present-day Japanese but also with ancient groups such as Yayoi- and Kofun-period individuals. Such consistently low values point to a deep divergence between the Jomon and all subsequent populations in the Japanese archipelago, likely because continental migrants—genetically distinct from the Jomon—entered the archipelago after the Yayoi period and admixed with Jomon-descended groups, as indicated by the ADMIXTURE analysis.

### 3.4 Differential Contributions of Jomon Groups to Present-day Japanese

To assess which Jomon groups may have contributed differentially to present-day Japanese ancestry, we computed *f*_4_-statistics in the form *f*_4_(Mbuti, Japanese; X, Y), where X and Y represent different Jomon groups, with the chronologically older group assigned as X (Figure 5). The *f*_4_ analysis revealed significant differences in genetic affinity between present-day Japanese and temporally distinct Jomon groups. Comparisons between the Initial Jomon individuals from Iyai and other Jomon individuals yielded consistently positive *f*_4_ values, with the strongest signals observed for *f*_4_(Mbuti, Japanese; Iyai, Todoroki) = 0.000993 (Z = 3.001) and *f*_4_(Mbuti, Japanese; Iyai, Chidorikubo) = 0.000982 (Z = 3.199). These results suggest that present-day Japanese individuals share more genetic drift with certain Early and Middle Jomon individuals, particularly Todoroki and Chidorikubo, than with Initial Jomon individuals. Comparisons with Mitsusawa and Funadomari showed the same directional trend, although with weaker statistical support (Z = 2.676 and 1.861, respectively). In contrast, all pairwise comparisons among non-Iyai Jomon individuals resulted in |Z| < 1, indicating similar levels of affinity with present-day Japanese. As a comparative reference, we also evaluated affinities with present-day Koreans, which showed no detectable variation among Jomon groups (Table S5).

**Figure 5.**
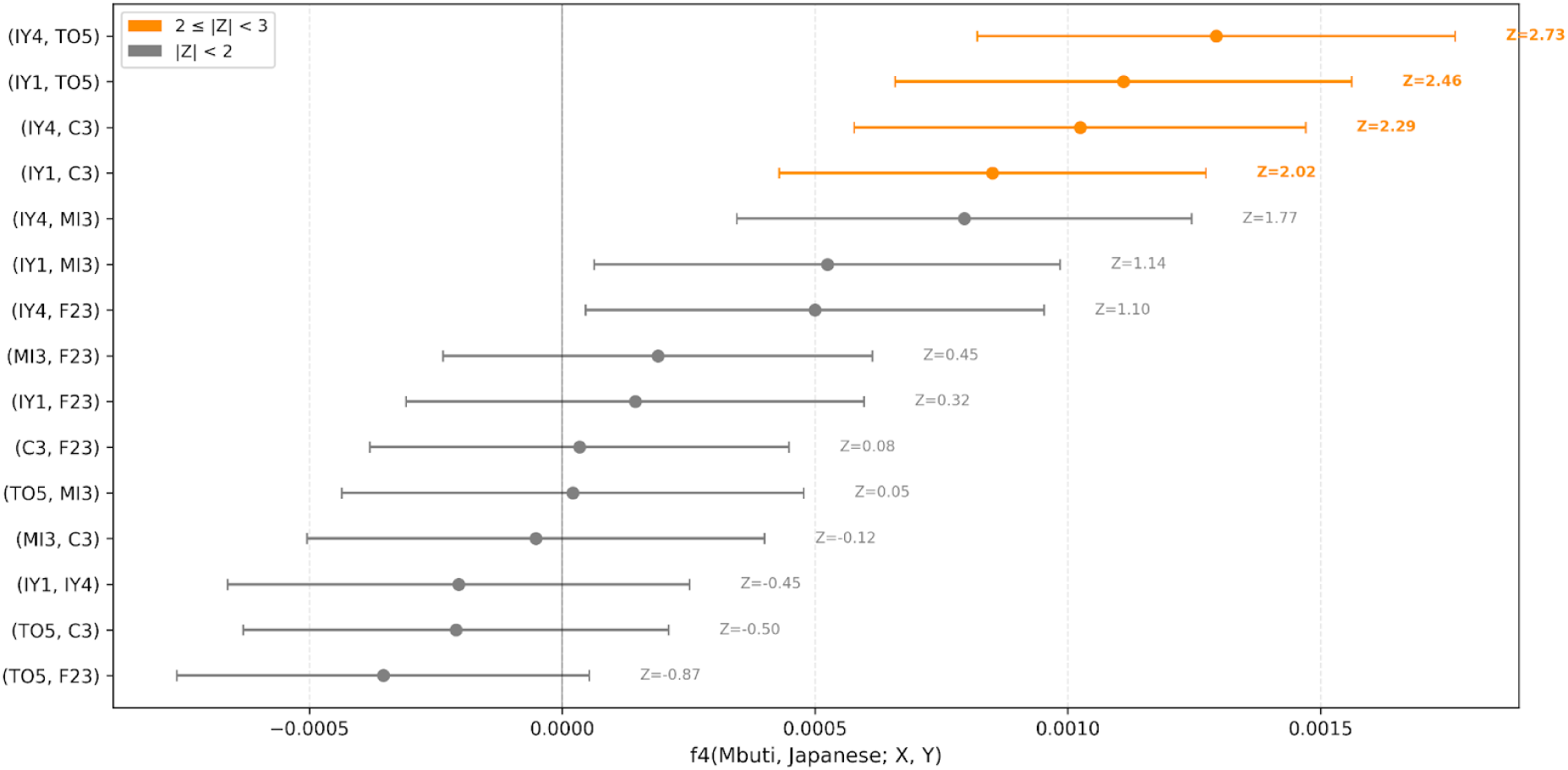
*f*_4_-statistics indicate differential genetic affinities between present-day Japanese and Jomon groups. Forest plot of *f*_4_ statistics of the form f_4_(Mbuti, Japanese; X, Y), where X and Y are different Jomon groups. The plot includes all 15 possible pairwise comparisons derived from the six Jomon individuals: IY1, IY4, TO5, MI3, C3, and F23. Squares represent f_4_ values with 95% confidence intervals. Results with 2 ≤ |Z| < 3 are shown in orange, whereas those with |Z| < 2 are shown in gray. Positive values indicate that present-day Japanese share more genetic drift with group Y than with X.

To quantify Jomon ancestry proportions in present-day mainland Japanese, we applied *f*_4_-ratio analysis based on a model of admixture between a Jomon source and a continental East Asian proxy, represented by present-day Koreans, using Mbuti as the outgroup (Figure 6). When IY1 was used as the Jomon source, present-day mainland Japanese exhibited 14.1% Jomon ancestry relative to Korean (SE = 0.0438, Z = 19.6), while IY4 yielded 15.1% (SE = 0.0425, Z = 20.0). In contrast, using other Jomon individuals consistently produced higher estimates, ranging from 17.2% to 19.4%. These values are substantially higher than previous estimates of ∼10% based on the Higashimyo Jomon (Adachi et al., 2021) and ∼13% based on the Funadomari Jomon (F23) (Kanzawa-Kiriyama et al., 2019).

**Figure 6.**
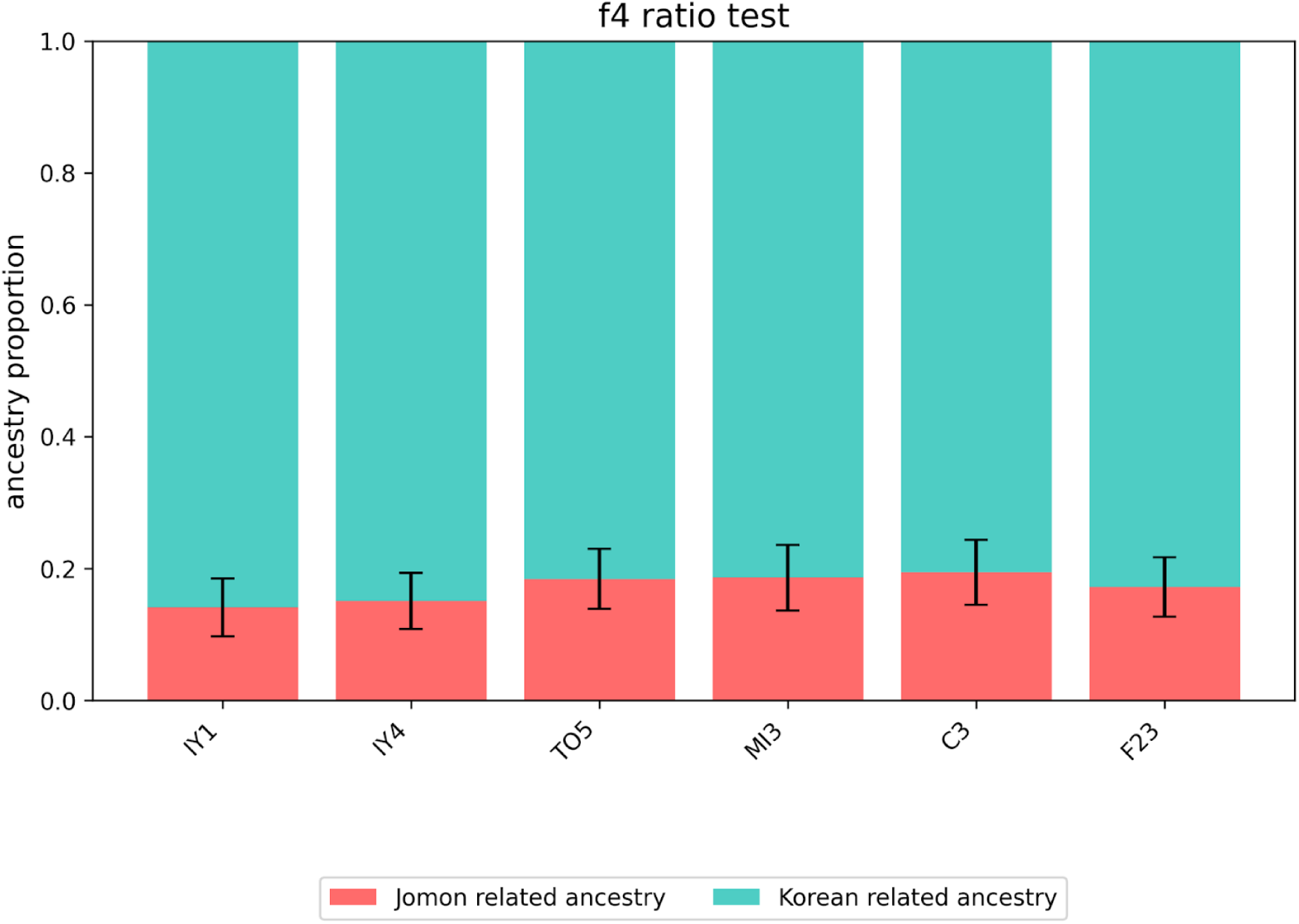
Estimates of Jomon ancestry in present-day Japanese from *f*_4_-ratio analysis. Bar plots showing the proportions of Jomon (red) and continental East Asian ancestry (turquoise) in present-day Japanese, estimated using *f*_4_-ratio analysis with present-day Koreans as a proxy for continental East Asian ancestry. Labels below each bar indicate individual IDs: IY1 = Iyai1, IY4 = Iyai4, TO5 = Todoroki5, MI3 = Mitsusawa3. C3 = Chidorikubo3, and F23 = Funadomari23. Error bars represent standard errors computed via block jackknife method.

### 3.5 Genetic Contributions of Jomon Individuals to the 47 Prefectures of Japan

To examine both the genetic contribution of Jomon individuals and the influence of population structure in present-day Japanese, we calculated outgroup *f*_3_ statistics between each Jomon individual and populations from all 47 prefectures (Figure 7). All Jomon individuals exhibited a similar geographic pattern, showing the lowest affinities with populations from the Kinki region and gradually higher values toward peripheral regions, particularly Kyushu and Tohoku.

**Figure 7.**
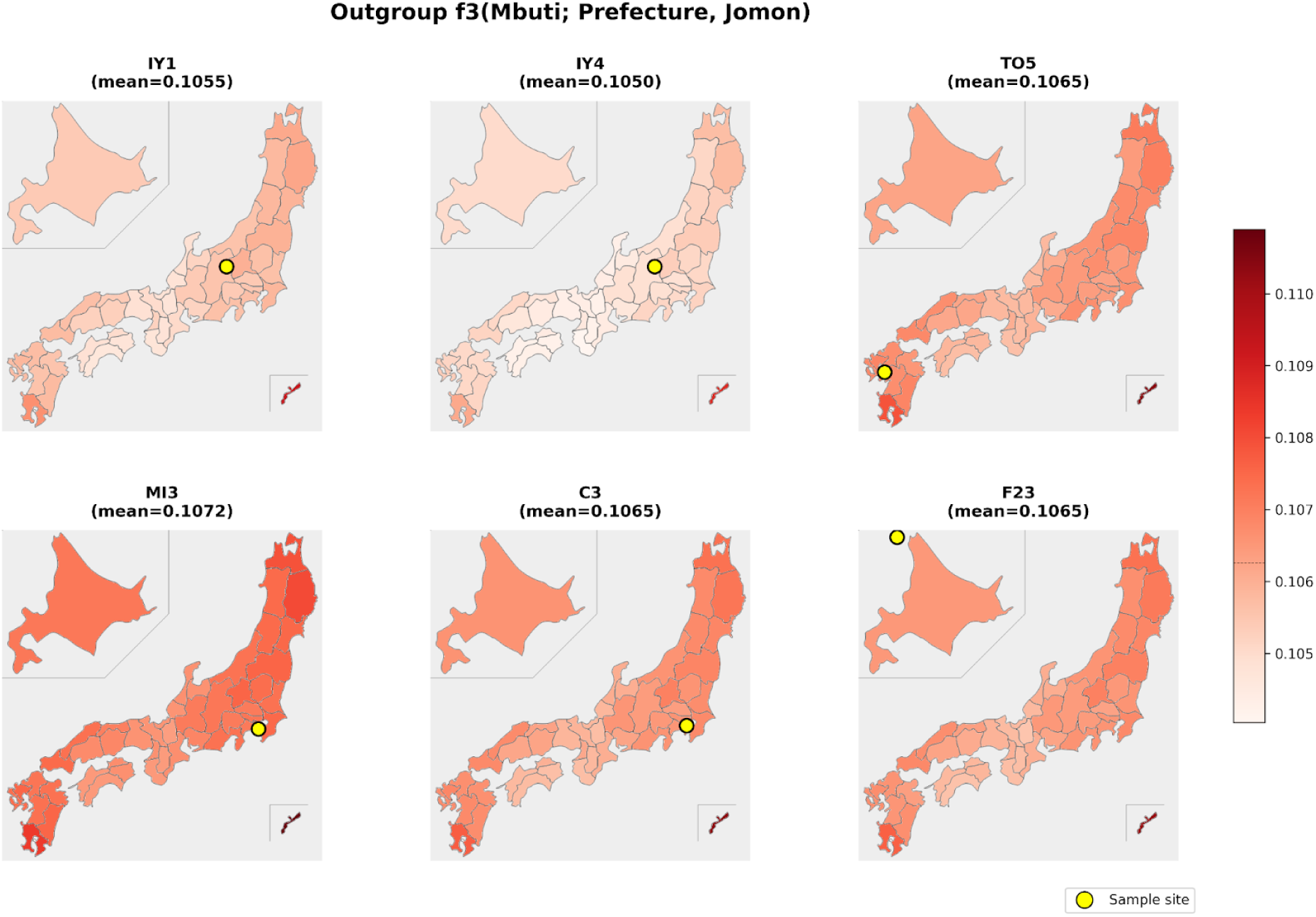
Geographic distribution of genetic affinity between Jomon individuals and present-day Japanese prefectural populations. Maps of Japan showing outgroup *f*_3_ values between individuals from 47 prefectures and six Jomon individuals: IY1 (Iyai1), IY4 (Iyai4), TO5 (Todoroki5), MI3 (Mitsusawa3), C3 (Chidorikubo3), and F23 (Funadomari23). Outgroup *f*_3_-statistics, *f*_3_(Mbuti; prefecture, Jomon), were calculated to assess genetic affinities between Jomon individuals and present-day populations from the 47 prefectures of Japan. Colors represent *f*_3_ values, with darker red indicating higher genetic affinity. The locations of Jomon individuals are indicated by yellow circles on the maps.

### 3.6 Branching order of Jomon groups

To examine phylogenetic relationships among Jomon groups, we performed TreeMix analysis incorporating present-day East Asian populations (Figure 8). In TreeMix, the drift parameter represents the amount of genetic drift that has accumulated along each branch since divergence and is roughly proportional to the number of elapsed generations relative to the effective population size. Branch lengths in the inferred tree are scaled according to this parameter. The Jomon groups exhibited longer branches than other populations (i.e., larger drift parameters), likely reflecting both their extended isolation since divergence and their relatively small effective population sizes.

**Figure 8.**
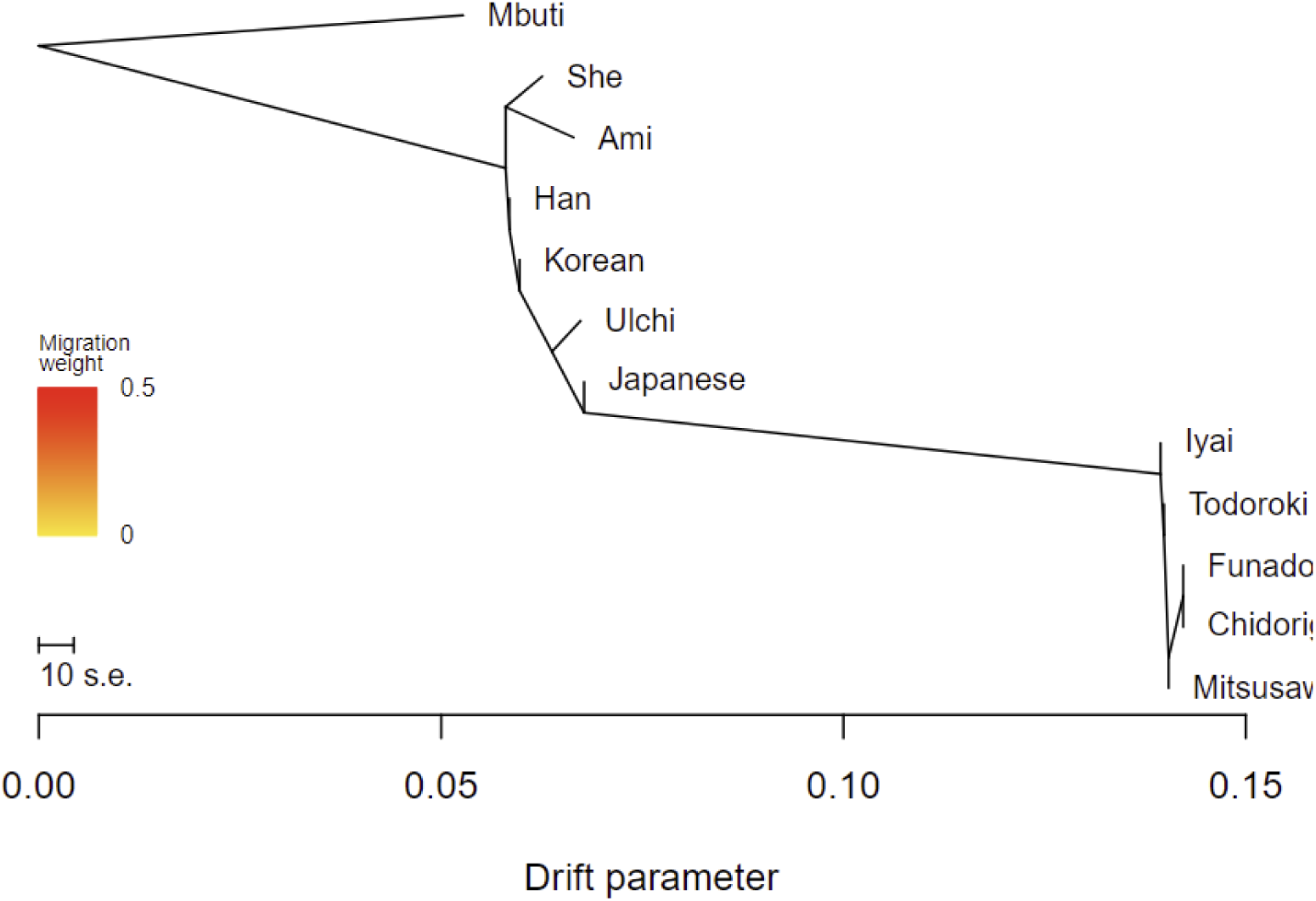
Phylogenetic relationships among Jomon and East Asian populations inferred by TreeMix. Maximum likelihood tree generated using TreeMix, including Jomon and present-day East Asian populations. The Iyai group (Initial Jomon) form the earliest branch within the Jomon clade, followed by chronologically ordered divergence of Todoroki (Early), Chidorikubo and Mitsusawa (Middle), and Funadomari (Late). The scale bar represents ten times the standard error of the entries in the covariance matrix.

The inferred tree topology within the Jomon clade indicated that the Iyai group (Initial Jomon) represented the earliest-diverging branch, followed by the sequential divergence of Todoroki (Early Jomon), Chidorikubo and Mitsusawa (Middle Jomon), and Funadomari (Late Jomon) (Figure 8). This branching pattern closely mirrors both the archaeological periods assigned to the samples and their 14C ages (Table 1), and it is consistent with the temporal genetic differentiation observed in the *f*-statistics analyses. However, because TreeMix roots the tree based on the included reference populations and does not account for ancient population structure outside the specified model, it incorrectly inferred the Jomon lineage as branching off from present-day Japanese in this analysis. This is a methodological artifact and should not be interpreted as reflecting the true ancestral relationship between these groups.

**Table 1.** Jomon individuals primarily analyzed in this study, with associated archaeological information, 14C ages, morphological sex assessments, and references.

| Sample | Site (Prefecture) | Archaeological period | 14C age (yBP)* | Morphological sex | Reference for morphological sex |
| --- | --- | --- | --- | --- | --- |
| IY1 | Gunma | Initial Jomon | 7413 ± 28 | Female | Kondo et al. (2018) |
| IY4 | Gunma | Initial Jomon | 7341 ± 35 | Undetermined | This study |
| IY15 | Gunma | Initial Jomon | 7438 ± 23 | Undetermined | This study |
| IY18 | Gunma | Initial Jomon | 7544 ± 26 | Male | This study |
| TO5 | Kumamoto | Early Jomon | 5340 ± 25 | Female | This study |
| C3 | Tokyo | Middle Jomon | 4593 ± 21 | Undetermined | This study |
| MI3 | Kanagawa | Middle Jomon | 4100 ± 21 | Female | This study |
| F23 | Hokkaido | Late Jomon | 3960–3550 | Female | Kanzawa-Kiriyama et al. (2019) |

\*Radiocarbon dates for IY4, IY15, IY18, TO5, C3, and MI3 were previously obtained; only 14C ages (yBP) are shown here. Calibrated dates and full measurement details will be presented in a separate publication.

### 3.7 Polygenic score–based inference of phenotypic traits in Jomon individuals

To assess the genetic characteristics associated with various quantitative traits, we calculated polygenic scores (PSs) for Jomon individuals using effect sizes from genome-wide association studies in Japanese (http://jenger.riken.jp/result) (Figure 9). Because PS analyses require reliable diploid genotype calls at each SNP, we restricted these analyses to the six high-coverage individuals. All scores were evaluated relative to the mean PS values of present-day Japanese. Given that the number of Jomon genomes analyzed (n = 6) is much smaller than the present-day Japanese sample set, no formal statistical tests were performed. Instead, we focus on traits for which all six Jomon individuals consistently showed PS values above or below the present-day Japanese mean.

**Figure 9.**
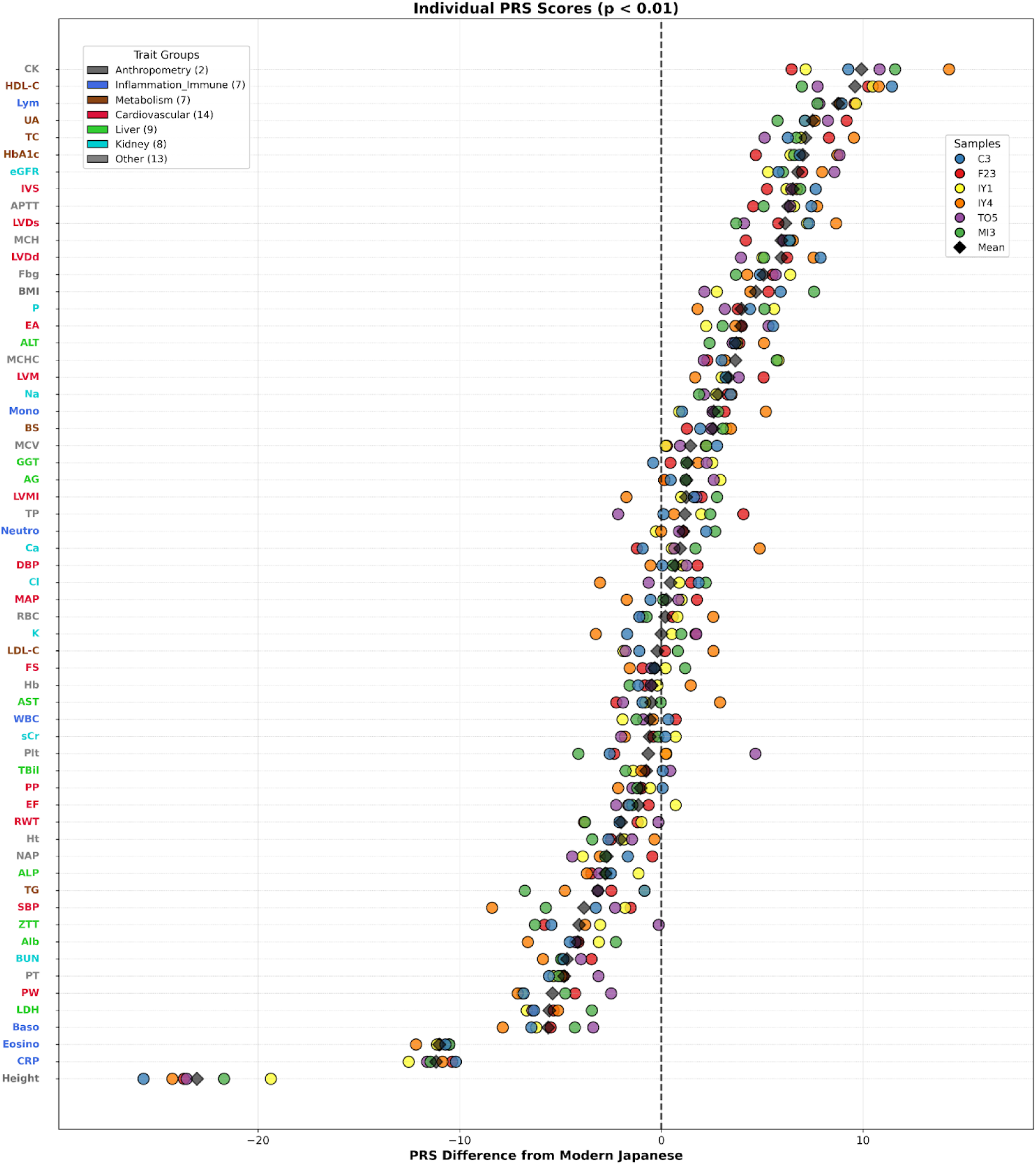
Polygenic score analysis reveals phenotypic differentiation among Jomon individuals. Forest plot showing polygenic risk score (PRS) differences between individual Jomon samples and present-day Japanese across traits significant at P < 0.01. Each point represents one of six Jomon individuals (C3, F23, IY1, IY4, TO5, MI3) color-coded as indicated in the legend. The x-axis shows PRS differences from present-day Japanese, with positive values indicating higher scores and negative values indicating lower scores compared to the present-day Japanese baseline (set at zero, indicated by the vertical dashed line). Traits are color-coded by functional category on the y-axis: anthropometry (gray), inflammation/immune (blue), metabolism (dark blue), cardiovascular (red), liver (green), kidney (turquoise), and others (light blue).

The PS analyses indicated that Jomon individuals differed from present-day Japanese in their genetic predispositions for multiple quantitative traits. In terms of anthropometric traits, they showed lower PSs for height but higher PSs for body mass index (BMI), consistent with differences in body size and shape. Several metabolic traits displayed systematic shifts: polygenic scores for HbA1c, total cholesterol (TC), uric acid (UA), blood sugar (BS), and high-density lipoprotein cholesterol (HDL-C) were elevated, indicating a genetic predisposition to higher levels of these traits in Jomon individuals. In contrast, immunological traits such as basophil counts, eosinophil counts, and C-reactive protein (CRP) showed reduced scores, suggesting a genetic predisposition to lower levels of these traits. Finally, the cardiovascular trait left ventricular dimensions (LVDs) revealed a predisposition toward larger LVDs, suggesting a tendency toward cardiac dilation.

## 4. DISCUSSION

Our analysis of eight high-coverage Jomon genomes spanning approximately 6,000 years reveals new insights into the genetic structure and evolution of hunter-gatherer populations in the Japanese archipelago. The tight clustering of all Jomon individuals in PCA plot (Figure 2) and their derivation from a single ancestral component in ADMIXTURE analysis (Figure 3) demonstrate substantial genetic homogeneity across both temporal and geographic scales. This genetic stability persisted despite the diverse ecological zones occupied by Jomon populations—from the subtropical environments of southern Kyushu to the temperate forests of central Japan and the subarctic conditions of Hokkaido. Such long-term genetic continuity over more than 13,000 years is exceptional among documented hunter-gatherer populations and likely reflects the archipelago’s geographic isolation following post-glacial sea level rise between 20,000 and 15,000 years ago (Lambeck et al., 2014; Skoglund &Mathieson, 2018).

Previous studies have suggested that the effective population size of the Jomon was around 1,000 individuals (Gakuhari et al., 2020; Cooke et al., 2021), and our TreeMix analysis supports this estimate. This demographic constraint would have limited genetic diversity while promoting genetic drift, resulting in the unique genetic signature that distinguishes Jomon populations from all other East Asian groups. The absence of detectable gene flow from continental populations in our ADMIXTURE and *f*-statistics analyses further supports a scenario of profound isolation. This contrasts markedly with contemporary hunter-gatherer populations in mainland Asia, such as those in the Amur River basin, who show evidence of repeated admixture with neighboring agricultural populations (Siska et al., 2017; Sikora et al., 2019).

Despite overall genetic homogeneity, our analyses of high-coverage Jomon genomes reveal clear temporal stratification, and together with regional variation, highlight heterogeneity within the Jomon gene pool. The Initial Jomon individuals from Iyai consistently show lower affinity with present-day Japanese compared to other Jomon groups (Figure 5), indicating their position as an outgroup within the broader Jomon cluster. This pattern, also confirmed by TreeMix analysis, identifies Iyai as the earliest branch within the Jomon clade (Figure 8). This observation is in agreement with a recent study reporting that the Initial Jomon individual from Shikoku diverged from later Jomon populations (Jeong et al., 2023). The significant *f*_4_-statistics demonstrate that present-day Japanese share more genetic components with TO5 and C3 than with other Jomon groups (Figure 5), suggesting that the genetic contribution to present-day Japanese was not uniform across Jomon populations from different regions. A key achievement of this study is the provision of high-quality Jomon genomes genetically closer to present-day Japanese than F23 (Kanzawa-Kiriyama et al., 2019), which has served as the reference Jomon genome in most previous studies. Using TO5 and C3, we show that the proportion of Jomon ancestry in mainland Japanese (∼20%) (Figure 6) is substantially higher than the ∼13% estimated from F23. Because our analyses rely on high-coverage genomes, individuals yielding higher ancestry estimates can be regarded as genetically closer to present-day Japanese. Looking ahead, TO5 and C3 are expected to serve as new reference Jomon genomes that will advance future research on population history from the Jomon period to the present.

The geographic distribution of genetic affinity between Jomon individuals and the present-day Japanese populations across the 47 prefectures provides important insights into the genetic continuity between regional Jomon groups and their corresponding present-day populations, and also sheds light on demographic processes that occurred after the Jomon period. All Jomon individuals display a shared geographic pattern, showing the lowest *f*_3_ values with populations from the Kinki region and progressively higher values toward the Tohoku and Kyushu regions (Figure 7). This overall pattern, observed consistently across all Jomon individuals analyzed in this study, is largely consistent with findings from a previous study (Watanabe and Ohashi, 2023). However, among the Jomon groups, the Iyai individuals consistently exhibit weaker genetic affinities with the present-day Japanese populations from all prefectures, likely reflecting their older chronological age. If the Jomon had constituted a highly structured population, present-day regional variation among the 47 prefectures would be expected to show closer genetic continuity with local Jomon groups. However, such continuity was not observed, suggesting that the Jomon as a whole did not form a strongly structured population. Instead, the regional differences observed among present-day Japanese are more likely to reflect demographic processes that unfolded after the Jomon period, particularly those following admixture with continental migrants. The concentration of continental ancestry in the Kinki region—the historical center of state formation during the Yayoi and Kofun periods—further supports archaeological models of agricultural expansion and political centralization emanating from central Japan (Barnes, 2015; Mizoguchi, 2013).

The high coverage of most Jomon genomes (>30×) enabled accurate diploid genotype calling, which in turn allowed us to examine genetic predispositions related to anthropometric, metabolic, immunological, and cardiovascular traits through PS analyses at the individual level (Figure 8). Importantly, Jomon individuals show reduced PSs for height, consistent with morphological studies that have long demonstrated their shorter stature (Hiramoto, 1972; Kaifu, 1992; Wada et al., 2000; Cox et al., 2019). This concordance indicates that our PS-based framework robustly captures genetic predispositions for quantitative traits from ancient genomes and offers a reliable basis for exploring other traits. Overall, Jomon individuals appeared to carry distinctive genetic predispositions for quantitative traits compared with present-day Japanese.

Extending this analysis, we found clear patterns of differentiation in metabolic and immunological traits among Jomon individuals, which align with long-standing expectations from evolutionary theory while providing the first direct ancient genomic evidence for these patterns. Jomon individuals show elevated PSs for BMI and several metabolic traits associated with efficient energy storage, including HbA1c, TC, BS, and HDL-C. These predispositions would have been advantageous for hunter-gatherer populations experiencing seasonal resource fluctuation and periodic scarcity, consistent with the thrifty genotype hypothesis (Neel, 1962). In contrast, Jomon individuals show reduced PSs for immunological traits such as basophil counts, eosinophil counts, and CRP—patterns indicative of limited selective pressure from parasitic and infectious diseases in the small, mobile hunter-gatherer societies characteristic of the Jomon population (Fumagalli et al., 2011; Quintana-Murci, 2019). Taken together with the finding that more than 80% of the genetic ancestry of present-day Japanese derives from continental migrants, these results imply that continental ancestors carried substantially higher PSs for immune-related traits, likely reflecting adaptation to the pathogen-rich environments characteristic of sedentary, wet-rice agricultural communities. These observations are broadly consistent with earlier ancestry-based work that inferred similar tendencies across metabolic and immunological traits using Jomon-derived variants estimated from present-day Japanese genomes (Watanabe and Ohashi, 2023). That study also proposed adaptive interpretations, including the thrifty genotype hypothesis and heightened pathogen pressures among continental agriculturalists. However, those inferences were based solely on population-averaged PSs for the Jomon ancestry. By contrast, our direct analysis of multiple ancient individuals enables us to evaluate not only the mean but also the variance of PSs within the Jomon population, revealing consistently low inter-individual differences across most traits examined (Figure 9). Such resolution cannot be achieved from population-averaged PSs alone. Moreover, whereas Watanabe and Ohashi (2023) inferred higher triglyceride (TG) PSs for the Jomon, our direct ancient-genome analyses instead indicate lower values, demonstrating the improved resolution afforded by individual-level ancient genomes and highlighting limitations inherent to indirect inference based solely on present-day Japanese genomes.

Finally, our PS analyses indicate a genetic predisposition for larger LVDs in the Jomon population, suggesting a tendency toward cardiac dilation. Such morphology may have been advantageous in a hunter-gatherer context by supporting sustained physical activity and endurance, as increased ventricular size can enhance cardiac output during prolonged exertion (Pluim et al., 2000). Larger ventricular dimensions may also have facilitated the acute hemodynamic demands of hunting or transporting heavy loads. Although left ventricular dilation is associated with elevated risks of chronic heart failure and arrhythmia (Yancy et al., 2013), these late-onset complications may have had limited opportunity to manifest given the shorter life expectancy of Jomon people (approximately 30–40 years; Mizoguchi, 2013; Habu, 2004). Thus, traits that could be deleterious today may have been largely inconsequential within their ecological and demographic context.

Our findings provide an integrated view of Jomon population history, revealing deep genetic continuity across space and time alongside clear temporal stratification within the Jomon lineage. By combining high-coverage ancient genomes with population genomic and polygenic score analyses, we refine estimates of Jomon ancestry in mainland Japanese and identify characteristic biological predispositions shaped by long-term hunter-gatherer adaptations. These ancient genetic features, together with the demographic processes involving admixture between Jomon people and continental migrants in the Japanese archipelago, continue to influence phenotypic variation and disease susceptibility in present-day Japanese. Our study demonstrates how ancient genomic data can illuminate past adaptations and their enduring impact on present-day human populations.

## Code availability

The study did not involve the development of new code. All software and scripts used are publicly available and referenced in the Methods section.

## Author contributions

F.M., S.U., and J.O. conceived and supervised the study. Y.T., M.M., and T.M. provided the archaeological materials and associated contextual information. O.K. and A.S. performed sex estimation based on skeletal morphology. F.M. and S.U. conducted the ancient DNA laboratory work. M.K., F.M., K.I., S.A., and J.K. processed the NGS data, performed read alignment, and carried out variant calling. M.Y. provided preliminary chronological assessments of the skeletal remains. Y.W. and M.I. assembled the prefecture-level allele frequency dataset. M.K. and I.N. curated the archaeological and genetic datasets. M.K. performed the statistical analyses and drafted the original manuscript. F.M. wrote the sections describing the experimental and NGS methodology, and J.O. provided revisions and editorial guidance. K.K. and L.W. contributed to discussions and interpretation of the results. All authors reviewed and approved the final version of the manuscript.

