## Supplementary Information for "High-Coverage Jomon Genomes Provide Insights into Population Structure and Genetic Traits of Ancient Japanese Hunter-Gatherers"

and Welfare, Niigata, Japan

6. The University Museum, The University of Tokyo, Tokyo, Japan

7. School of Medicine, Hangzhou Normal University, Hangzhou, China

8. Institute for the Study of Ancient Civilizations and Cultural Resources, Kanazawa University

9. Medical Genome Center, National Research Institute for Child Health and Development, Tokyo, Japan

10. Department of Genetics, Albert Einstein College of Medicine, NY, USA

*To whom correspondence may be addressed:

Fuzuki Mizuno, Ph.D.

Department of Legal Medicine, Toho University School of Medicine, 5-21-16, Omori-Nishi, Ota-ku, Tokyo, 143-8540, Japan.

Jun Ohashi, Ph.D.

Department of Biological Sciences, Graduate School of Science, The University of Tokyo, 7-3-1, Hongo, Bunkyo-ku, Tokyo, 113-0033, Japan.

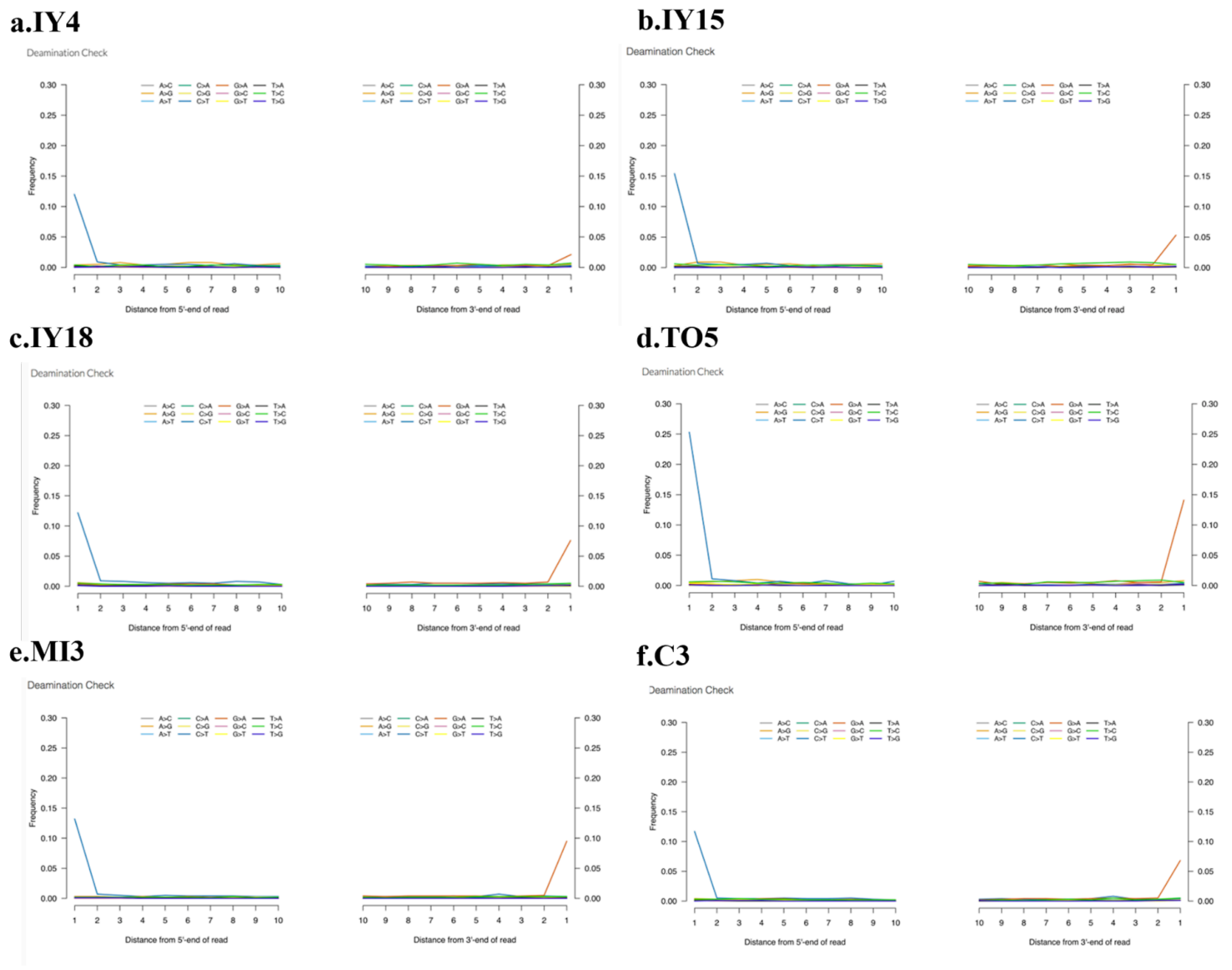


**Figure S1. The deamination pattern of the double stranded library of Jomon individuals**

The mismatch frequency is relative to the reference as a function of read position; C to T in blue and G to A in red. Panels a–f correspond to the individuals indicated in the figure.

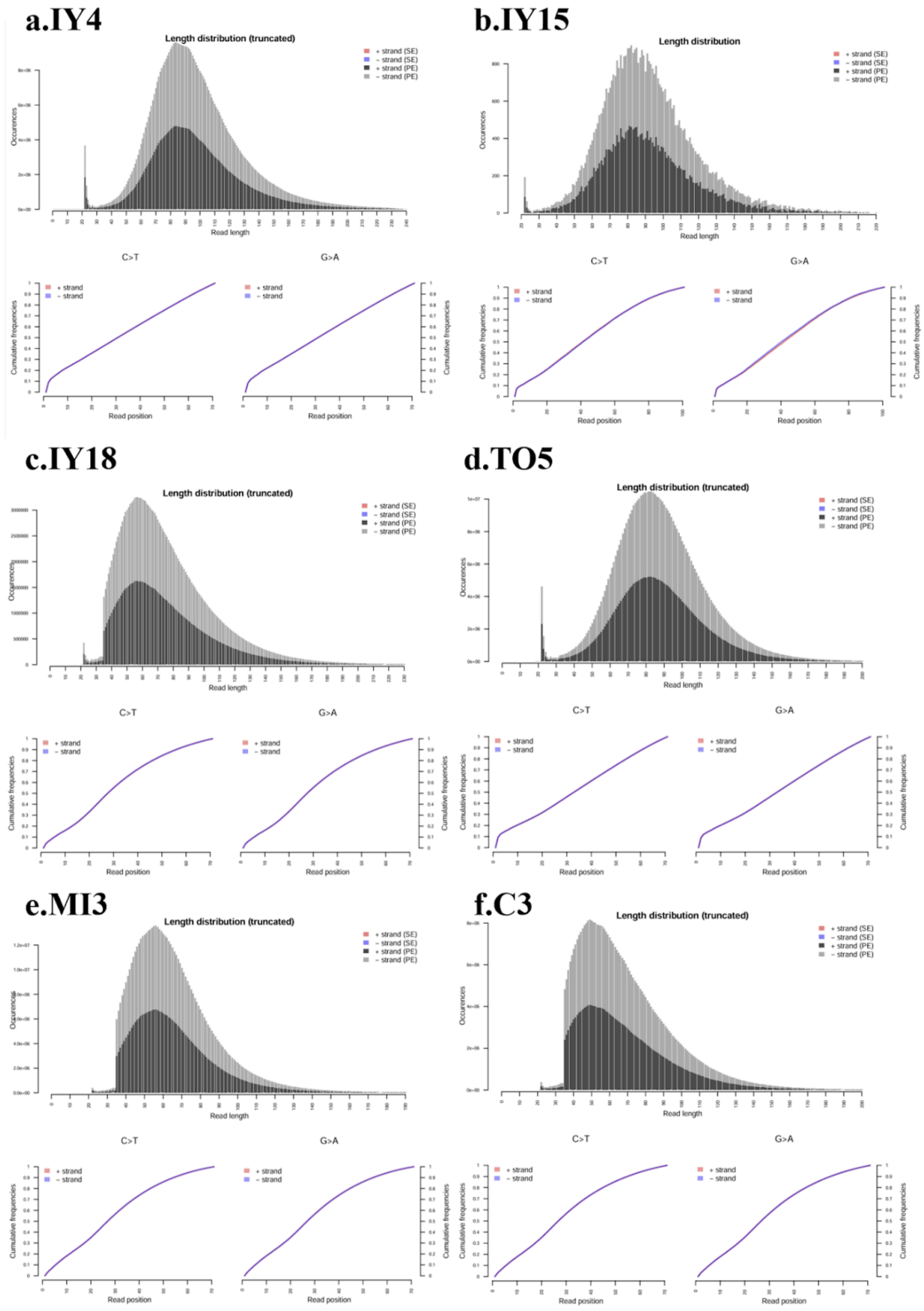


**Figure S2. Fragment-length distribution and terminal damage profiles**

Fragment-length distributions (top) and cumulative frequencies of 5′ C→T and 3′ G→A (bottom; left and right), estimated by mapDamage. Panels a–f correspond to the individuals indicated in the figure.

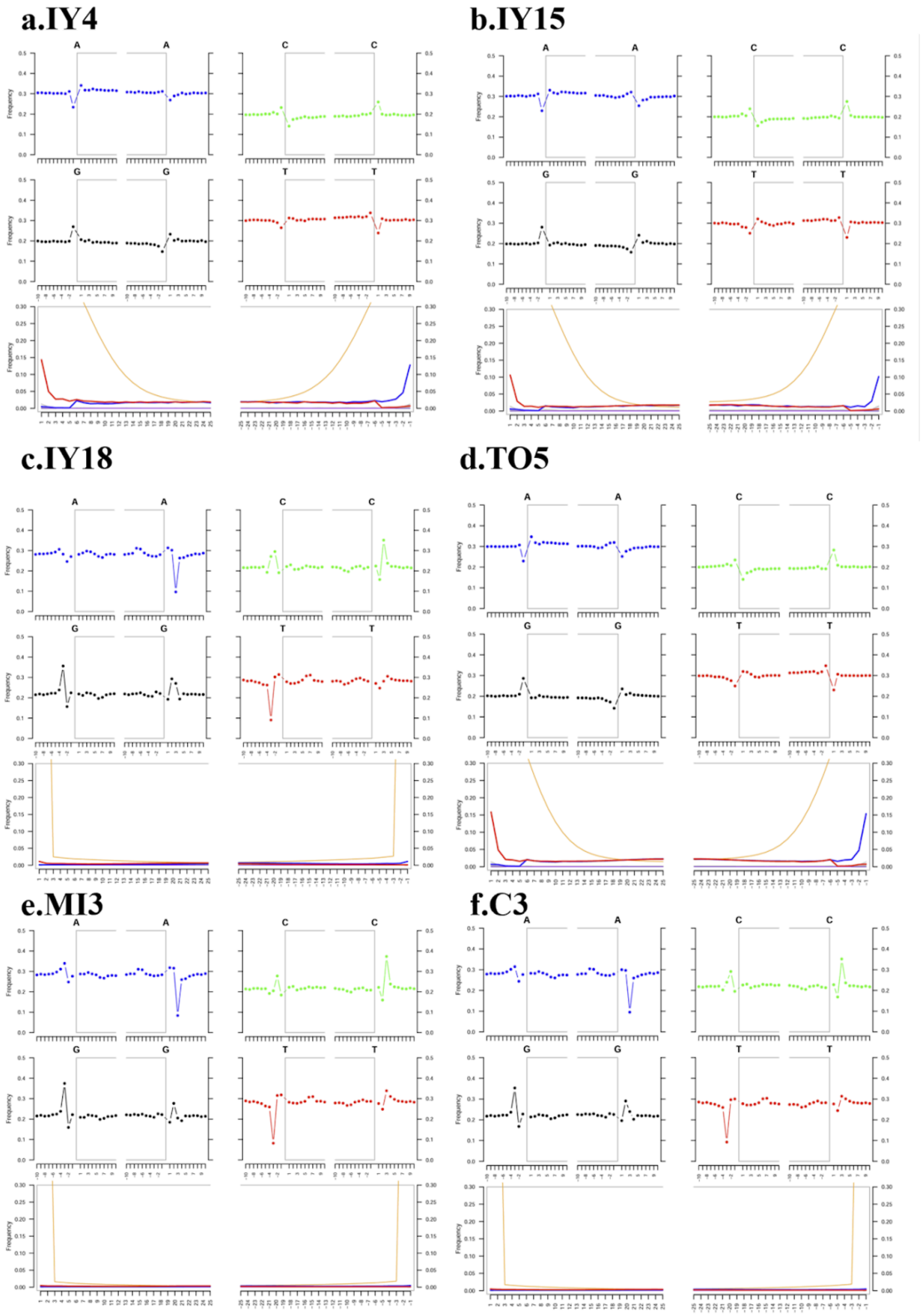


**Figure S3. Misincorporation patterns at read termini**

Fragmisincorporation patterns at read termini inferred by mapDamage. For each individual (see panel labels), the upper panels show position-dependent substitution frequencies for each reference base (A/C/G/T) (left: 5′ end; right: 3′ end; gray boxes indicate internal read positions/background). The lower panels show position-dependent frequencies of C→T at the 5′ end (left) and G→A at the 3′ end (right), plotted for the + strand (red), − strand (blue), and both strands combined (purple). The orange line represents a smoothed/fit curve to the observed data.

**Table S1. Genetic sex determination of ancient Japanese individuals**

| Sample | X_reads | Y_reads | Total_XY | RY_index | Inferred sex |
|---|---|---|---|---|---|
| IY1 (Iyai1) | 114949649 | 332771 | 115282420 | 0.002886 | Female |
| IY4 (Iyai4) | 34798403 | 6833529 | 41631932 | 0.164141 | Male |
| IY15 (Iyai15) | 52303790 | 9991263 | 62295053 | 0.160386 | Male |
| IY18 (Iyai18) | 5403855 | 1302611 | 6706466 | 0.194232 | Male |
| TO5 (Todoroki5) | 86118485 | 234307 | 86352792 | 0.002713 | Female |
| MI3 (Mitsusawa3) | 65236923 | 2113896 | 67350819 | 0.031386 | Undetermined |
| C3 (Chidorikubo3) | 43265241 | 1370083 | 44635324 | 0.030695 | Undetermined |
| F23 (Funadomari23) | 66795661 | 97781 | 66893442 | 0.001461 | Female |

Sex was determined based on the RY index (Y chromosome reads / total sex chromosome reads) following Skoglund et al. (2013). Samples with RY < 0.016 were classified as female, RY > 0.075 as male, and intermediate values as undetermined.

**Table S2. X- and Y-chromosomal read counts obtained from targeted whole-genome in-solution enrichment sequencing of MI3 and C3**

| Individual ID | X_reads | Y_reads | Inferred sex |
|---|---|---|---|
| MI3 (Mitsusawa3) | 217,969 | 1,473 | Female |
| C3 (Chidorikubo3) | 201,846 | 1,405 | Female |

Read counts mapping to the X and Y chromosomes for individuals MI3 and C3, obtained after double-stranded (DS) library preparation, whole-genome in-solution enrichment, and sequencing on an Illumina MiSeq platform.

**Table S3. $f_4$-ratio test for Jomon individuals**

| A | B | X | C | Outgroup | alpha | 1-alpha | SE | Z |
|---|---|---|---|---|---|---|---|---|
| Dai | Korean | Japanese | C3 | Mbuti | 0.80619 | 0.19381 | 0.049345 | 16.338 |
| Dai | Korean | Japanese | F23 | Mbuti | 0.827952 | 0.172048 | 0.045233 | 18.304 |
| Dai | Korean | Japanese | IY1 | Mbuti | 0.858647 | 0.141353 | 0.043824 | 19.593 |
| Dai | Korean | Japanese | IY4 | Mbuti | 0.848846 | 0.151154 | 0.042543 | 19.953 |
| Dai | Korean | Japanese | MI3 | Mbuti | 0.81379 | 0.18621 | 0.049579 | 16.414 |
| Dai | Korean | Japanese | TO5 | Mbuti | 0.815693 | 0.184307 | 0.045479 | 17.936 |

**Table S4.** ***f*₄-statistics testing differences in genetic affinity to present-day Japanese among Jomon individuals**

| A | B | C | D | f4 | Z | nBABA | nABBA | nSNP |
|---|---|---|---|---|---|---|---|---|
| Mbuti | Japanese | C3 | F23 | 0.000034 | 0.082 | 17721 | 17711 | 293629 |
| Mbuti | Japanese | IY1 | C3 | 0.000851 | 2.018 | 17766 | 17524 | 284564 |
| Mbuti | Japanese | IY1 | F23 | 0.000144 | 0.318 | 17701 | 17660 | 289767 |
| Mbuti | Japanese | IY1 | IY4 | -0.00021 | -0.449 | 17509 | 17568 | 284093 |
| Mbuti | Japanese | IY1 | MI3 | 0.000525 | 1.138 | 17963 | 17812 | 288374 |
| Mbuti | Japanese | IY1 | TO5 | 0.00111 | 2.46 | 17538 | 17226 | 281322 |
| Mbuti | Japanese | IY4 | C3 | 0.001024 | 2.293 | 16885 | 16607 | 271877 |
| Mbuti | Japanese | IY4 | F23 | 0.0005 | 1.103 | 16730 | 16593 | 274452 |
| Mbuti | Japanese | IY4 | MI3 | 0.000795 | 1.767 | 17086 | 16866 | 276670 |
| Mbuti | Japanese | IY4 | TO5 | 0.001293 | 2.735 | 16759 | 16410 | 269807 |
| Mbuti | Japanese | MI3 | C3 | -0.000053 | -0.116 | 18021 | 18036 | 296618 |
| Mbuti | Japanese | MI3 | F23 | 0.000188 | 0.444 | 17965 | 17908 | 298043 |
| Mbuti | Japanese | TO5 | C3 | -0.00021 | -0.499 | 16605 | 16663 | 271743 |
| Mbuti | Japanese | TO5 | F23 | -0.00035 | -0.868 | 16542 | 16640 | 274907 |
| Mbuti | Japanese | TO5 | MI3 | 0.000021 | 0.047 | 16895 | 16889 | 276084 |

**Table S5.** ***$f_4$*****-statistics testing differences in genetic affinity to present-day Koreans among Jomon individuals**

| A | B | C | D | f4 | Z | nBABA | nABBA | nSNP |
|---|---|---|---|---|---|---|---|---|
| Mbuti | Korean | C3 | F23 | 0.000106 | 0.201 | 17084 | 17054 | 286254 |
| Mbuti | Korean | IY1 | C3 | 0.000309 | 0.583 | 17086 | 17000 | 278354 |
| Mbuti | Korean | IY1 | F23 | -0.00028 | -0.505 | 17052 | 17131 | 283412 |
| Mbuti | Korean | IY1 | MI3 | 0.000389 | 0.682 | 17357 | 17248 | 281977 |
| Mbuti | Korean | IY1 | TO5 | 0.001049 | 1.917 | 16931 | 16643 | 274958 |
| Mbuti | Korean | IY4 | C3 | 0.000458 | 0.869 | 16263 | 16141 | 265938 |
| Mbuti | Korean | IY4 | F23 | 0.000002 | 0.004 | 16130 | 16129 | 268448 |
| Mbuti | Korean | IY4 | MI3 | 0.000521 | 0.939 | 16489 | 16348 | 270593 |
| Mbuti | Korean | IY4 | TO5 | 0.000892 | 1.545 | 16159 | 15923 | 263727 |
| Mbuti | Korean | MI3 | C3 | -0.000072 | -0.136 | 17366 | 17387 | 289068 |
| Mbuti | Korean | MI3 | F23 | 0.000019 | 0.036 | 17285 | 17279 | 290535 |
| Mbuti | Korean | TO5 | C3 | -0.0003 | -0.573 | 16045 | 16125 | 265727 |
| Mbuti | Korean | TO5 | F23 | -0.0005 | -0.982 | 15966 | 16101 | 268815 |
| Mbuti | Korean | TO5 | MI3 | -0.00013 | -0.245 | 16291 | 16327 | 269907 |

**Table S6. Traits and GWAS References used to compute PSs in Jomon individuals**

| Traits | Reference |
|---|---|
| Body mass index (BMI) | Akiyama et al., 2017 |
| Height | Akiyama et al., 2019 |
| Albumin/globulin ratio (AG) | Kanai et al., 2018 |
| Alkaline phosphatase (ALP) | Kanai et al., 2018 |
| Alanine aminotransferase (ALT) | Kanai et al., 2018 |
| Activated partial thromboplastin time (APTT) | Kanai et al., 2018 |
| Aspartate aminotransferase(AST) | Kanai et al., 2018 |
| Albumin (Alb) | Kanai et al., 2018 |
| Blood sugar (BS) | Kanai et al., 2018 |
| Blood urea nitrogen (BUN) | Kanai et al., 2018 |
| Basophil count (Baso) | Kanai et al., 2018 |
| Creatine kinase (CK) | Kanai et al., 2018 |
| C-reactive protein (CRP) | Kanai et al., 2018 |
| Calcium (Ca) | Kanai et al., 2018 |
| Chloride (Cl) | Kanai et al., 2018 |
| Diastolic blood pressure (DBP) | Kanai et al., 2018 |
| E/A ratio (EA) | Kanai et al., 2018 |
| Ejection fraction (EF) | Kanai et al., 2018 |
| Eosinophil count (Eosino) | Kanai et al., 2018 |
| Fractional shortening (FS) | Kanai et al., 2018 |
| Fibrinogen (Fbg) | Kanai et al., 2018 |
| γ-glutamyl transferase (GGT) | Kanai et al., 2018 |
| High-density-lipoprotein cholesterol (HDL-C) | Kanai et al., 2018 |
| Hemoglobin (Hb) | Kanai et al., 2018 |
| Hemoglobin A1c (HbA1c) | Kanai et al., 2018 |
| Hematocrit (Ht) | Kanai et al., 2018 |
| Interventricular septum thickness (IVS) | Kanai et al., 2018 |
| Potassium (K) | Kanai et al., 2018 |
| Lactate dehydrogenase (LDH) | Kanai et al., 2018 |
| Low-density-lipoprotein cholesterol (LDL-C) | Kanai et al., 2018 |
| Left ventricular internal dimension in diastole (LVDd) | Kanai et al., 2018 |
| Left ventricular internal dimension in systole (LVDs) | Kanai et al., 2018 |
| Left ventricular mass (LVM) | Kanai et al., 2018 |
| Left ventricular mass index (LVMI) | Kanai et al., 2018 |

| | |
|---|---|
| Lymphocyte count (Lym) | Kanai et al., 2018 |
| Mean arterial pressure (MAP) | Kanai et al., 2018 |
| Mean corpuscular hemoglobin (MCH) | Kanai et al., 2018 |
| Mean corpuscular hemoglobin concentration (MCHC) | Kanai et al., 2018 |
| Mean corpuscular volume (MCV) | Kanai et al., 2018 |
| Monocyte count (Mono) | Kanai et al., 2018 |
| Non-albumin protein (NAP) | Kanai et al., 2018 |
| Sodium (Na) | Kanai et al., 2018 |
| Neutrophil count (Neutro) | Kanai et al., 2018 |
| Phosphorus (P) | Kanai et al., 2018 |
| Pulse pressure (PP) | Kanai et al., 2018 |
| Prothrombin time (PT) | Kanai et al., 2018 |
| Posterior wall thickness (PW) | Kanai et al., 2018 |
| Platelet count (Plt) | Kanai et al., 2018 |
| Red blood cell count (RBC) | Kanai et al., 2018 |
| Relative wall thickness (RWT) | Kanai et al., 2018 |
| Systolic blood pressure (SBP) | Kanai et al., 2018 |
| Total bilirubin (TBil) | Kanai et al., 2018 |
| Total cholesterol (TC) | Kanai et al., 2018 |
| Triglyceride (TG) | Kanai et al., 2018 |
| Total protein (TP) | Kanai et al., 2018 |
| Uric acid (UA) | Kanai et al., 2018 |
| White blood cell count (WBC) | Kanai et al., 2018 |
| Zinc sulfate turbidity test (ZTT) | Kanai et al., 2018 |
| Estimated glomerular filtration rate (eGFR) | Kanai et al., 2018 |
| Serum creatinine (sCr) | Kanai et al., 2018 |
